# Divergent *Hox* cluster collinearity in horned beetles reveals adult head patterning function of *labial*

**DOI:** 10.1101/2025.02.07.637090

**Authors:** Erica M. Nadolski, Isabel G. Manley, Sukhmani Gill, Armin P. Moczek

## Abstract

*Hox* genes play critical roles in specifying the regionalization of the body axis across metazoa, with the exception of the anterior dorsal head of bilaterian animals, which instead is instructed by a deeply conserved set of non-*Hox* regulators. The anterior dorsal head is also a hot spot of evolutionary diversification, raising the question as to the developmental-genetic underpinnings of such innovation. Onthophagine dung beetles develop evolutionarily novel and highly diversified horns on the dorsal head used as weapons during intrasexual conflicts. Preliminary RNAseq data unexpectedly documented *Hox* gene expression in the dorsal head of premetamorphic onthophagine larvae. Motivated by this observation, we aimed to (i) investigate the genomic content and arrangement of the *Hox* cluster across three onthophagine species, and (ii) assess expression patterns and (iii) potential functions of the anterior Hox genes *labial, proboscipedia*, and *Deformed* in patterning the adult beetle head and cephalic horns. We document an unexpected, derived *Hox* cluster configuration in the *Onthophagus sagittarius* genome, a species with apomorphic cephalic horn morphology. Yet despite this genomic rearrangement, embryonic expression patterns of *labial* and *proboscipedia* as well as the adult segment patterning functions of *proboscipedia* and *Deformed* were found to be conserved. In contrast, *labial* RNAi revealed an adult head patterning function outside horn-forming regions previously undescribed for any insect. Lastly, we show that electrosurgical ablation of the presumptive larval *labial*-expressing head region phenocopies this conspicuous adult *labial* RNAi defect. We discuss the implications of these data for current models of insect head development and diversification.

## Introduction

The majority of the arthropod body is composed of serially homologous segments that differentiate as instructed by *Hox* gene expression along the anteroposterior axis during early embryonic development (Lewis 1978, Carroll 1995). A major exception to this rule is the anterior-most region of the embryo, the ocular/protocerebral region (for simplicity, ocular region), which is patterned by a set of non-*Hox* regulators and is considered to be non-segmental in origin (Rogers & Kaufman 1997, Abzhanov & Kaufman 1999, Posnien *et al*. 2011, Zattara *et al*. 2016). Those segments that are patterned by Hox genes undergo homeotic (identity) transformations when the corresponding *Hox* genes are functionally manipulated (Hughes & Kaufman 2000, Brown *et al*. 2002). These data led to the historical proposal of the *straight-face model* of head development, wherein the post-embryonic head is made up of segments consisting of both ventral and dorsal components, matching the basic architecture of their thoracic and abdominal counterparts (Figure 1A, top). However, studies in a variety of insect orders have documented that dorsal head structures remain unaffected after *Hox* manipulations (Rogers *et al*. 2002, Posnien & Bucher 2010, Smith & Jockusch 2014). A later theory termed the *bend-and-zipper* model incorporates this observation by positing that the dorsal head instead originates via the upfolding, posterior bending, and fusing of the *Hox*-free and non-segmental antero-lateral embryonic primordia, with gnathal *Hox*-expressing segments contributing only to the ventral head (Posnien & Bucher 2010, Posnien *et al*. 2010, Figure 1A-B). This newer model successfully incorporated the bulk of existing data on insect head morphogenesis during the transition from the embryonic to juvenile stage but is agnostic to morphogenetic dynamics during the metamorphic transition from the juvenile to adult stage.

**Figure 1.**
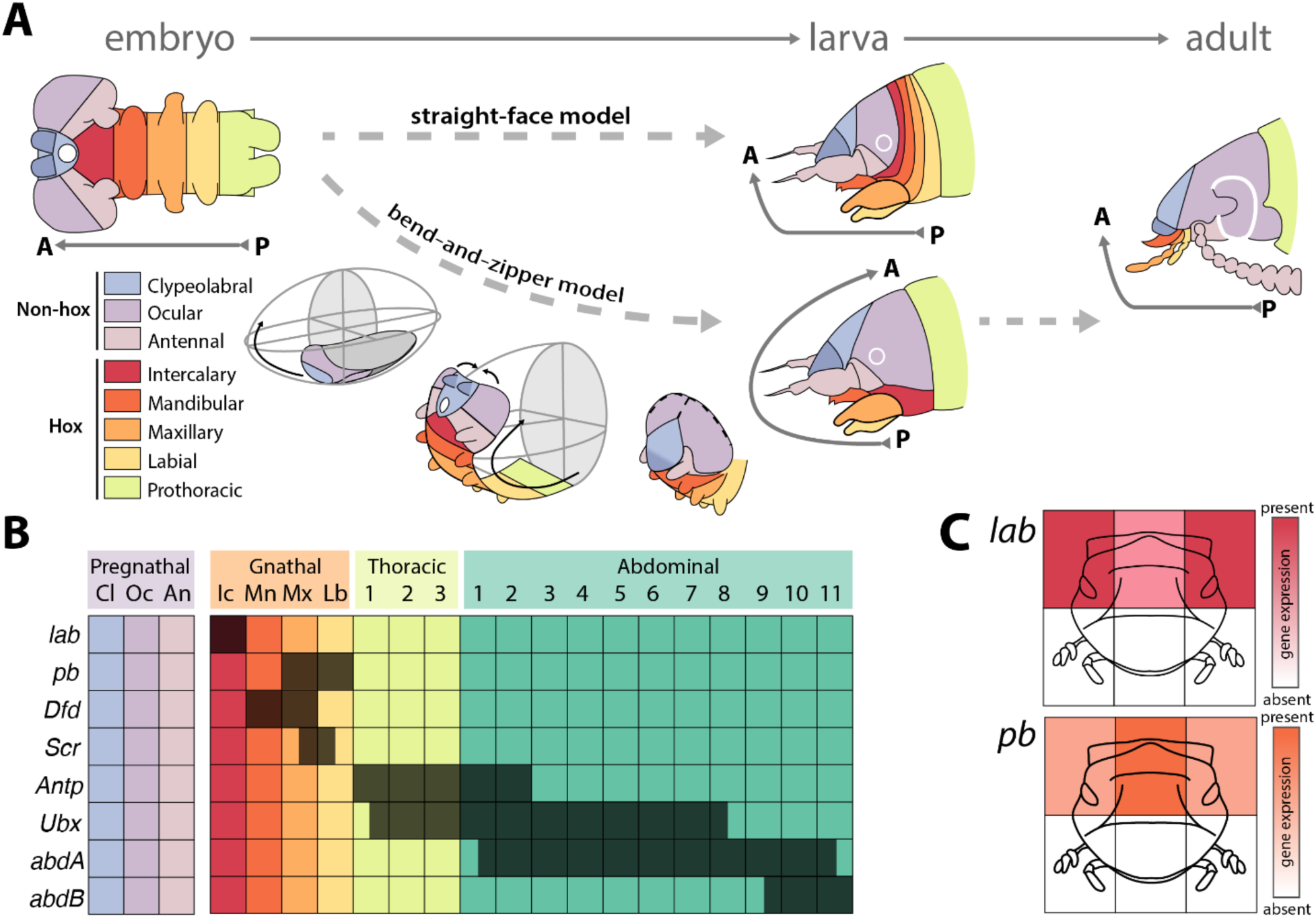
Current models of insect head morphogenesis and *Hox* gene expression data across beetles. To contrast the traditional straight-face model of insect head development, Posnien *et al*. (2010) proposed the bend-and-zipper model, wherein the anterior end of the flat early embryo folds upward and then fuses along the dorsal midline to meet the prothoracic segment (A). This model accounts for *Tribolium* data documenting that parental RNAi of *Hox* genes affects only ventral, mouthpart-bearing regions or the lateral gena but leaves the dorsal head of larvae unaffected. Additional work in *Tribolium* recapitulated these mouthpart transformations in adults after larval *Hox* RNAi (A, right). Expression patterns of each *Hox* gene along the segmented *Tribolium* embryo are shown in (B, dark boxes). Transcriptomic data from sections of the dorsal head epithelium of pre-pupal *O. taurus* document expression of *lab* and *pb* in the posterior half of the dorsal head (C). Segment abbreviations: Cl – clypeolabral; Oc – ocular; An – Antennal; Ic – Intercalary; Mn– mandibular; Mx – maxillary; Lb – labial; [Diagrams in A are based on Posnien *et al*. 2010 (Fig1C, 2B-D), Posnien & Bucher 2010 (Fig 5M), and Smith & Jockusch 2014; data in B from Brown *et al*. 2002, data in C from Linz & Moczek 2020].

In addition to being a presumptively *Hox*-free region, the dorsal head of insects is also a hot spot of evolutionary innovation, having given rise to *e.g.* the weevil rostrum, the stalks of stalk-eyed flies, and the cephalic horns of scarab beetles. Yet, how morphological novelties such as horns and stalks may originate and then integrate within deeply conserved trait complexes such as the dorsal head without compromising ancestral functions remains largely unclear. Horned dung beetles, in particular, are famous for their extraordinarily diversified dorsal cephalic horns, a textbook example of evolutionary novelties (Emlen *et al*. 2005). In the horned dung beetle genus *Onthophagus* alone, cephalic horns differ in number, positioning, and relative size as a function of species, sex, and sometimes simply adult size. Ancestral state reconstructions suggest, however, that this enormous diversity was initiated when male *Onthophagus* evolved the ability to form one or multiple horns in the posterior head specifically (Emlen *et al*. 2005). Ablation-based fate mapping studies on two species exhibiting this ancestral pattern of horn growth – *Onthophagus taurus* and *Digitonthophagus gazella –* determined that paired posterior head horns are positioned roughly along the boundary between the clypeolabral and ocular regions, both of which form from within the non-segmental anterior-most embryonic head (Busey *et al*. 2016). A closely related species, *O. sagittarius*, exhibits a derived horn morphology in the dorsal posterior head, with males *repressing* posterior horn formation (Kijimoto *et al*. 2012) and females sporting a single, medial head horn. Fate mapping of the female horn also indicated a different regional origin, arising instead from fully within the boundaries of the clypeolabral region (Busey *et al*. 2016).

A subset of onthophagine species also possess independently evolved *prothoracic horns*, which recent studies have identified as partial serial homologs of the meso- and metathoracic wings of insects, as well as other dorsal and dorsolateral traits including the bilateral gin traps observed in the abdomen of *Tribolium* and *Tenebrio* pupae (Hu *et al*. 2019, Hu & Moczek 2021). One key piece of evidence in support of this conclusion derived from the observation that RNAi-mediated transcript depletion of *Sex combs reduced*, the *Hox* gene instructing prothorax identity, contributes to the transformation of the prothoracic horn of *Onthophagus* beetles into ectopic wings, a key requirement for establishing (serial) homology. In contrast, the evolutionary origins of head horns remain largely unclear, including the possibility of anterior *Hox* gene involvement. However, recent work utilizing head-region-specific RNAseq raised precisely this possibility. Specifically, a comparative transcriptomic study in *O. taurus* wherein lateral, medial, posterior, and anterior regions of the dorsal head epithelium were microdissected and sequenced as separate libraries found the two anterior-most *Hox* genes *labial* and *proboscipedia* to be upregulated in posterolateral head regions (Linz & Moczek 2020, Figure 1C). These data stand at odds with predictions made by the *bend-and-zipper* model as well as our general understanding of insect head evolution which instead assumed *Hox* patterning to be restricted to ventral portions of the head (Figure 1A).

Motivated by the documentation of *Hox* gene expression in the dorsal head of horned beetle larvae, we aimed to first investigate the genomic arrangement of the *Hox* cluster in three onthophagine species. We then assessed the expression patterns of the anterior *Hox* genes *labial* and *proboscipedia* using *in situ* Hybridization Chain Reaction (HCR), followed by a combination of larval RNAinterference (RNAi) and electrosurgical ablation approaches to investigate their possible function in patterning the adult beetle head and/or cephalic horns. More generally, we aimed to use these data to further our understanding of the segmental contributions to the adult beetle head and to reconcile the *bend-and-zipper* model of insect head development with the dorsal *Hox* expression data observed in horned dung beetles.

## Methods

### Assessing genomic content and arrangement of Hox clusters

Custom BLAST databases were generated for the *O. sagittarius*, *O. taurus,* and *D. gazella* proteomes (Davidson & Moczek 2024) using command-line BLAST+ (Altschul *et al*. 1990). *Drosophila melanogaster* Hox gene protein sequences were downloaded from Flybase (version FB2024_02, Öztürk-Çolak *et al*. 2024) as queries. The query sequences were used to search the beetle proteome databases to identify target protein sequences for the eight canonical insect Hox genes involved in regionalization of the anteroposterior body axis: *Labial, Proboscipedia, Deformed, Sex combs reduced, Antennapedia, Ultrabithorax, Abdominal-A* and *Abdominal-B*. We established an e-value cutoff of 1×10^−5^ for selecting the best hit from each list of potential targets and manually validated the top hits (Table 1) with the end-to-end sequence alignment tool MUSCLE (MUltiple Sequence Comparison by Log-Expectation, Madeira *et al*. 2024). Upon discovering an unexpected arrangement of the cluster in one genome, we wanted to secondarily validate this finding; to do so, we reconstructed the contigs generated during genome sequencing from the raw PacBio HiFi CCS long read sequencing data via *hifiasm v. 0.13* under default parameters (Cheng *et al*. 2021), which we then examined for the presence of the entire derived *Hox* cluster configuration on a single contig.

### Beetle husbandry

*Onthophagus sagittarius* individuals were collected near Kilcoy, Queensland, AU (-26.951, 152.605), and *O. taurus* individuals were collected near Chapel Hill, North Carolina, USA (35.936, -79.128). Both species were reared in lab colonies as described previously (Moczek & Nagy 2005). Reproductively active adults were transferred from the colonies into breeding containers and allowed to reproduce. After one week, eggs and eclosed larvae were transferred from their natal brood balls into twelve-well plates provisioned with cow dung from Marble Hill Farm, Bloomington, Indiana, USA, as described in Shafiei *et al*. (2001). Plates with larvae were kept at a 16:8h light/dark cycle throughout development, with *O. taurus* incubated at 24°C and *O. sagittarius* incubated at 28°C pre- and post-injection until eclosion. Eclosed experimental adults were sacrificed and preserved in 70% ethanol.

### Embryonic dissections and in situ hybridization chain reaction

We selected *O. sagittarius* embryos within three days of egg-laying for sample preparation. The embryos were dissected out of their eggshells in 8% formaldehyde (FA) solution and allowed to fix for 40 minutes at room temperature. After several washes with PBT, the tissue was dehydrated through washes in 25%, 50%, 75% and 100% methanol in PBST. The samples were kept in 100% methanol overnight at −20 °C, rehydrated through a series of washes in 75%, 50%, 25% methanol in PBST, and rinsed several times in PBST. Tissues were then subjected to a 5-minute proteinase K (10 μg/μl) digestion, rinsed several times in PBST, post-fixed in 3.7% FA for 20 min, and washed in PBT. The samples were subsequently processed using standard procedures of third generation *in situ* hybridization chain reaction (*in situ* HCR v3.0) (Choi *et al*. 2018). DAPI counterstain was also applied to the tissues to visualize nuclei. After several washes in sodium chloride sodium citrate buffer with 0.05% Triton-X, tissues were mounted on a glass slide in glycerol. A Leica Stellaris 8 confocal microscope equipped with a HC PL APO CS2 10x/0.40 dry objective was used to image whole-mount embryos using a white light laser (WLL: 440nm - 790nm) and a Diode 405nm laser, acousto-optical tunable filter, a pinhole of 1 AU, and the following detectors per fluorophore (target, fluorophore, detector, emission tuning/gain/offset): DAPI, HyD S1, 420-510nm/91/0; *Os-pb*, AlexaFluor488, HyD S1, 505-560nm/53/0; *Os-lab*, AlexaFluor647, HyD X3 660-780nm/98/0.

### Double-stranded RNA synthesis and injection for RNA interference

Fragments of each target gene for RNA interference (RNAi) were chosen by using a BLAST algorithm to query 250bp regions of each gene against the *O. taurus* or *O. sagittarius* transcriptome and selecting regions with zero off-target hits. Oligonucleotide constructs for each target gene and construct-specific primers were designed using the appropriate reference genome (Davidson and Moczek 2024) and ordered from Integrated DNA Technologies, Inc. Synthesis of double-stranded RNA (dsRNA) for gene knockdown via RNAi was performed using a protocol optimized for coleopteran larvae (Philip & Tomoyasu 2011). Briefly, for each gene, PCR was performed to anneal T7 RNA polymerase binding sequences to the ends of the gene fragment construct, generating the DNA template for dsRNA synthesis. The Qiagen QIAquick PCR Purification kit was used to purify these constructs. In vitro transcription to generate dsRNA from the DNA template was performed using an Ambion MEGAscript T7 kit, and each product was then purified using an Ambion MEGAclear kit with an ethanol precipitation step (Philip & Tomoyasu 2011).

Double-stranded RNA constructs for each target gene were diluted to 1 µg/µL with injection buffer (Philip & Tomoyasu 2011). A Hamilton brand syringe and small gauge removable needle (32 gauge) were used to inject a 3µl dose of dsRNA targeting a single gene through the abdominal cuticle into the hemolymph of third-instar larvae, to initiate whole-body RNAi. For experimental treatments targeting multiple genes simultaneously, injection mixture was diluted to a 1µg/µl concentration per target gene, with injection doses totaling 3ul. Control individuals were randomly selected from each round of developing larvae and injected with pure injection buffer. Previous work in dung beetles has shown that injection of pure buffer alone can serve as a suitable control for dsRNA injection, as neither buffer-injected nor nonsense RNA-injected adults have been documented to show any detectable phenotypic differences compared to wildtype adults (Moczek & Rose 2009, Simonnet & Moczek 2011, Linz *et al*. 2019).

### Fate mapping via electrosurgical ablation of larval head epithelium

A Hyfrecator 2000 electrosurgical unit (ConMed, Utica, NY) was fitted with a metal surgical tip (Epilation/Telangiectasia Needle Stealth ER coating 30 angle, 3/8, 714-S; ConMed) to execute precise ablations of the head epidermis through the larval cuticle in both species. Using protocols developed by Busey *et al*. (2016), we applied 10 Watts for 3 seconds (*O. taurus*) or 2 seconds (*O. sagittarius*), respectively, applications found to generate clearly distinguishable phenotypes without causing excess mortality. Three separate posterior dorsal head regions were evaluated, targeting cells on the right side of the head of each treated larvae while leaving the left side of the same head untreated to serve as a negative control. All animals were treated during the last larval instar stage prior to apolysis and epidermal proliferation occurring during the metamorphic transition.

### Phenotype assessment and photography

We analyzed the phenotypes of RNAi-injected, control buffer-injected, and voltage-ablated individuals after eclosion. After sacrificing adult animals, whole heads were dissected away from the body for clarity of viewing. For RNAi and control-injected animals, ventral mouthparts were dissected away from the head for clarity of viewing. Representative individuals from each sample group were photographed using a Leica MZ16 microscope with a PLANAPO 2.0x objective and a Leica S8APO microscope and a PixeLINK PL-D7912CU-T camera; multiple photos of each sample were taken across different planes of focus and overlaid using Adobe Photoshop.

## Results

Motivated by earlier work documenting unexpected Hox gene expression in the dorsal head of premetamorphic larvae (Linz and Moczek 2020), we aimed to investigate the genomic Hox cluster across multiple horned dung beetle species, as well as the potential functions of anterior Hox genes *labial, proboscipedia*, and *Deformed* in establishing the segmental boundaries of the adult beetle head and in patterning the cephalic horns contained therein. We first investigated the genomic arrangement and content of the Hox cluster in three species: *D. gazella* and *O. taurus*, which represent the ancestral cephalic horn morphology (Figure 2A-B), and *O. sagittarius*, which represents a derived cephalic horn morphology and a reversal in the sexual dimorphism of the posterior cephalic horn (Figure 2C). We then proceeded with subsequent functional analyses in *O. sagittarius* and one of the ancestral proxy species, *O. taurus*. Four salient results emerged.

**Figure 2.**
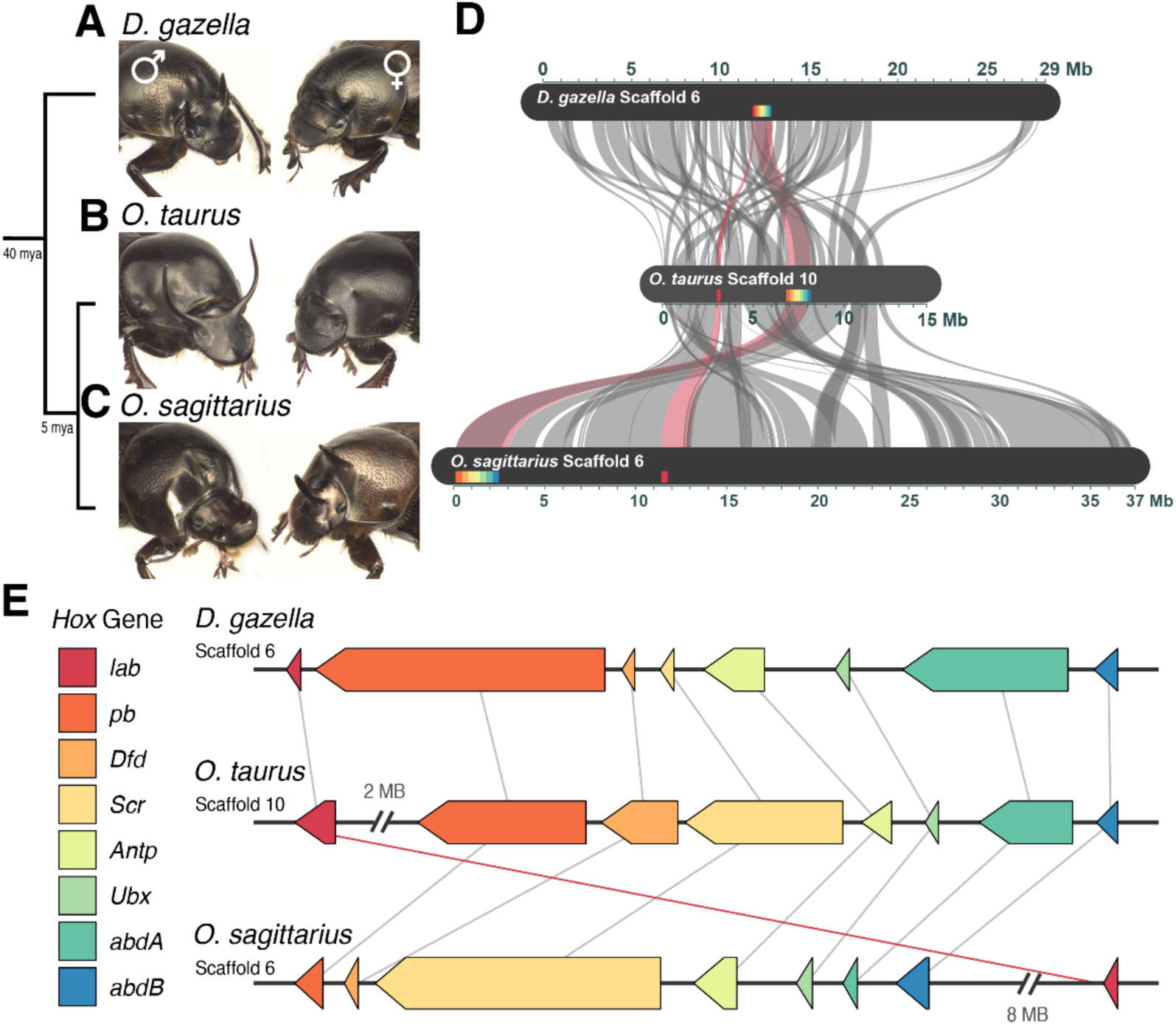
Genomic rearrangement of the *Hox* cluster in a dung beetle species with derived cephalic horn morphology. Onthophagine dung beetles possess evolutionarily novel and highly diversified head horns. (A) *D. gazella* males possess relatively modest paired posterior horns while females are hornless, whereas (B) *O. taurus* males possess greatly exaggerated paired posterior horns while females are, once again, hornless. (C) In contrast, *O. sagittarius* males are hornless in the posterior head, yet exhibit a novel pair of small anterior horns, while females develop a single medial posterior horn on the head (as well as a medial horn emanating from the prothorax not investigated in the present study). (D) In these three species, the scaffolds containing the *Hox* clusters are largely syntenic across their length, albeit showing a variety of internal translocations (grey bars), with synteny of Hox genes (colored boxes) highlighted as red bars. (E) Genomic collinearity of the *Hox* cluster is conserved in *D. gazella* and *O. taurus* but derived in *O. sagittarius* with the majority of the cluster translocated 8MB down to the 5’ end of the scaffold retaining its orientation, leaving *Os-lab* in its ancestral location more central to the scaffold (red line). Relative coding sequence lengths in E are represented accurately within but not across genomes.

### Genomic rearrangement of the Hox cluster in a species with derived horn morphology

We used a reciprocal BLAST approach to identify *Hox* gene orthologs in three horned dung beetle species for which sequenced genomes with chromosome level resolution are available: *O. sagittarius, O. taurus*, and *D. gazella*. Specifically, we targeted the eight canonical insect *Hox* genes that have retained their ancestral function in regional body identity specification during early embryogenesis: *lab, pb, Dfd, Scr, Antp, Ubx, abdA*, and *abdB*. This approach revealed an intact cluster with exactly one ortholog of each *Hox* gene in the expected 5’ to 3’ order in both *D. gazella* and *O. taurus* on scaffolds largely syntenic across their length (Figure 2D, Supplementary Table 1). In *O. taurus*, one large split nearing 2MB in length was detected separating *Ot-lab* from *Ot-pb* and the rest of the intact cluster. In the *O. sagittarius* genome, we also uncovered a set of exactly one ortholog per Hox gene on the scaffold largely syntenic to those containing the other dung beetle Hox clusters (Figure 2D). However, in this genome the canonical order of the cluster diverges (Fig 2E). Specifically, *Os-lab* is the only gene retaining a central location on the scaffold, while the rest of the cluster is translocated over 8MB 5’ on the same scaffold.

To validate this finding and rule out that this arrangement could be an artefact of the genome scaffolding process, we returned to the contigs generated through PacBio long read sequencing to assess cluster configuration directly. To do so, we generated a custom BLAST database comprised of the complete set of long-read contigs and performed BLAST searches for each *O. sagittarius* Hox query. In doing so, we confirmed that a single contig contained the entire modified Hox cluster in *O. sagittarius*, including the novel location of the *Os-pb – Os-abdB* cluster on the 5’ end of the scaffold, the 8 MB stretch intervening stretch, and the central location of *Os-lab* (Supplementary Table 2). This result supports the finding that this derived rearrangement is not an artefact of the genome assembly pipeline and instead reflects a novel translocation within the cluster potentially unique to the *O. sagittarius* genome.

### O. sagittarius maintains canonical Hox expression during embryogenesis

With the analyses described above indicating a rearrangement of the *Hox* cluster in *O. sagittarius*, we sought to assess whether *Os-labial* retained its canonical segment-restricted expression during embryonic development. To do so we used *in situ* HCR targeting *Os-lab* in embryos roughly 24 hours after egg laying (AEL) or roughly 72 hours AEL (Figure 3A) to capture a range of embryogenesis processes. We also targeted *Os-proboscipedia* in the same experiment as a positive control, since this gene has retained its close contiguity with the rest of the Hox cluster in the *O. sagittarius* genome. Staining 24h AEL embryos revealed *Os-lab* signal in the intercalary segment, i.e. the first of the serially homologous segments posterior to the ocular region, and thus a pattern typical for holometabolous insects (Figure 3B-D). Despite several attempts we were, however, unable to capture *Os-pb* staining at the same timepoint, potentially due to conserved temporal collinearity wherein *labial* expression appears prior to any subsequent *Hox* expression.

**Figure 3.**
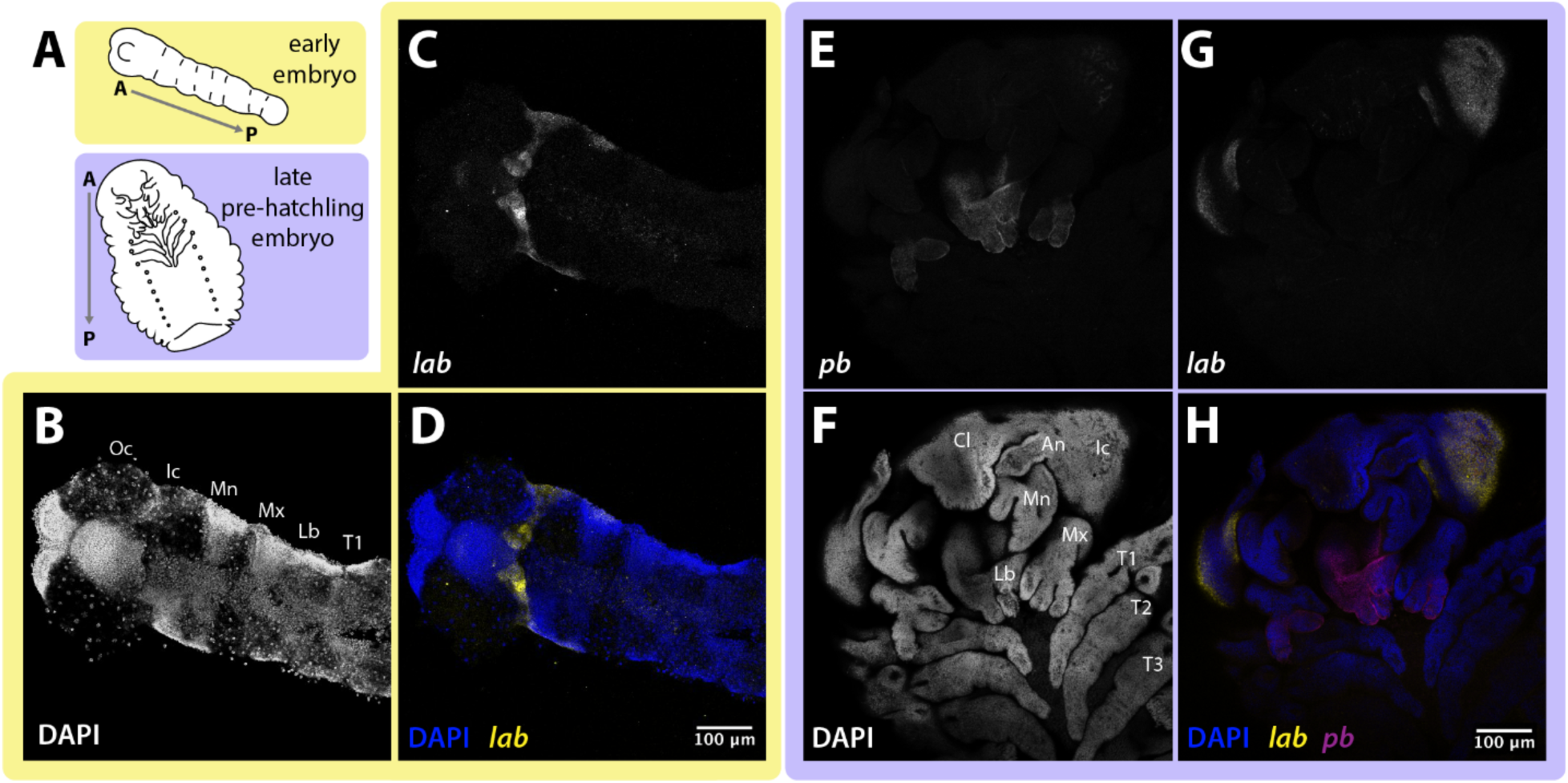
Canonical *lab* and *pb* expression patterns are maintained despite genomic rearrangement in *O. sagittarius.* Representative diagrams of developmental stages of embryos used for *in situ* HCR staining (A). Yellow outlining (B-D) indicates data from the early embryonic stage and purple outlining (E-H) indicates data from the late embryonic stage. In an early *O. sagittarius* embryo approximately 24h after egg laying (AEL) shown dorsally co-stained with DAPI (B), HCR staining of *lab* shows typical expression pattern in the first of the serially homologous segments in the body plan (C, D). In an older, pre-hatchling stage *O. sagittarius* embryo shown ventrally, HCR staining for *pb* (E) was performed alongside co-staining for DAPI (F) and *lab* (G). Staining at this stage confirms that *pb* is expressed in the expected segments which bear the maxillary and labial feeding appendages; *lab* expression at this stage appears bilaterally in the pre-hatchling head. Segment abbreviations: Cl – clypeolabral; Oc – ocular; An – antennal; Ic – intercalary; Mn – mandibular; Mx – maxillary; Lb – labial; T1-prothorax; T2 – mesothorax; T3 – metathorax.

In contrast, staining 72 hours AEL embryos revealed signal for both *Os-lab* and *Os-pb*. At this developmental stage, *Os-lab* expression was observed dorsal to the mouthparts and restricted to two distinct lateral rather than one contiguous head region (Figure 3F-H). *Os-pb* expression in turn was observed in the segments bearing the future maxillary and labial mouthparts, posterior to the presumed intercalary segment (Figure 3E). These results are thus consistent with a conservation of canonical segment patterning by *Os-lab* and *Os-pb*.

### Functional analysis of anterior Hox genes in horned dung beetles

Next, we analyzed the functions of *lab, pb*, and *Dfd* individually and in combination using larval RNAi in both *O. taurus* (conserved genomic Hox arrangement) and *O. sagittarius* (derived cluster configuration). We predicted that if anterior Hox gene functions are conserved in these species then RNAi targeting *pb* and *Dfd* – but not *lab –* should recapitulate homeotic transformations of ventral mouthparts well described across diverse insects (Hughes & Kaufman 2000, Smith & Jockusch 2014, Zhang *et al*. 2020). Furthermore, we predicted that RNAi-mediated transcript depletion of any of our three target genes should not affect *dorsal* head patterning in line with the assumption that this body region is not instructed by *Hox* gene input. Our findings match the first, but diverge from the second prediction.

Specifically, RNAi-mediated transcript depletion of *Os-lab* did not result in homeotic mouthpart transformations, matching predictions from previous work which established that the intercalary segment does not contribute to appendage formation in insects (Figure 4D). Also as predicted, *Os-pb* and *Ot-pb* RNAi animals exhibited a homeotic transformation of both maxillary palps and labial palps into legs, evidenced by the increased size of the proximal palpomeres, the decrease in size of the distal palpomeres, and the appearance of long sensory bristles on the proximal palpomeres, with particularly well-developed ectopic femurs borne by the maxillary palps (Figure 4E). *Os-Dfd* and *Ot-Dfd* RNAi animals also exhibited an expected knockdown phenotype in the maxilla, specifically a rounding of the everted corners of the base of the maxillary galea (Figure 4F), although no deformation was detected in the mandibles from *Dfd* RNAi alone, diverging from findings in *Tribolium* (Figure S1A-B).

**Figure 4.**
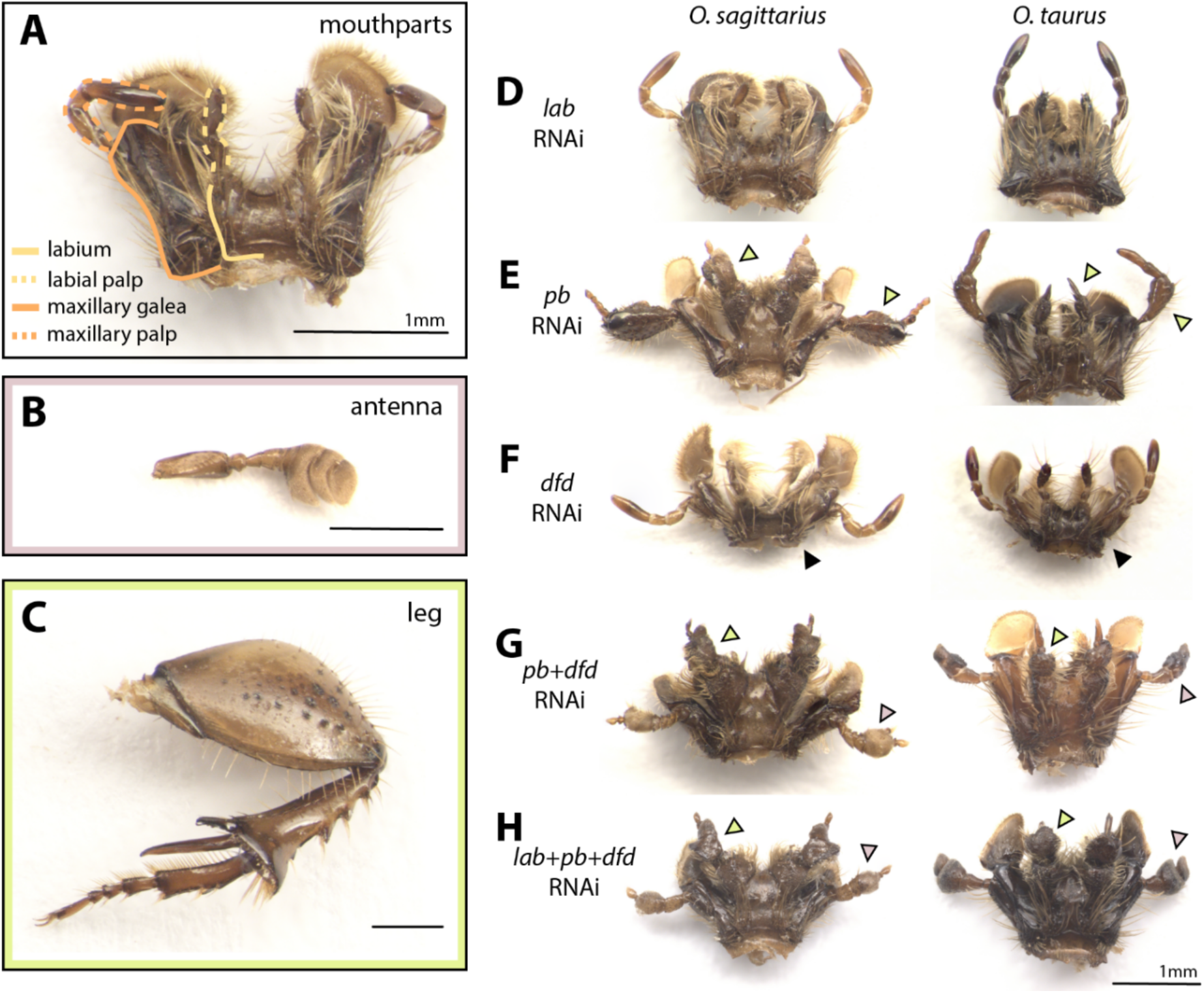
Homeotic transformations of adult mouthparts following larval *Hox* RNAi in dung beetles mirror those described for other beetle taxa. Buffer-injected control *O. sagittarius* illustrating wildtype appendages: mouthparts derived from the maxillary and labial segments (A) antenna (B), and mesothoracic leg (C). *Hox* gene knockdown via RNAi in *O. sagittarius* and *O. taurus* results in a range of homeotic transformations (D to H). *Os-lab* RNAi and *Ot-lab* RNAi did not result in changes to mouthpart morphology (D). *Os-pb* RNAi and *Ot-pb* RNAi resulted in transformation of the maxillary palps and labial palps to legs, indicated by the increased size of the proximal palpomeres, the decrease in size of the distal palpomeres, and the appearance of sensory bristles on the proximal palpomeres (E, green arrowheads). *Os-dfd* RNAi and *Ot-dfd* RNAi resulted in deformation of the maxillary galea, indicated by the smaller, rounded shape of the base (F, black arrowheads). Double RNAi of *pb* + *dfd* in *O. sagittarius* and *O. taurus* resulted in transformation of the maxillary palps to antennae (G, pink arrowheads) and the labial palps to legs (G, green arrowheads). Triple RNAi knockdown of *lab + pb* + *dfd* in *O. sagittarius* and *O. taurus* also resulted in transformation of the maxillary palps to antennae (H, pink arrowheads) and the labial palps to legs (H, green arrowheads).

Double knockdown of *pb* and *Dfd* in *O. sagittarius* and *O. taurus* resulted in a new set of homeotic transformations: labial palps transformed to express leg identity while maxillary palps acquired antennal identity, as evidenced by the enlargement of the medial and distal palpomeres and the appearance of densely packed, short sensory bristles. (Figure 4G). The double knockdown treatment led to a deformation of the mandibles, specifically the appearance of ectopic bristles on the molar region (Figure S1C). Further, triple knockdown of *lab+pb+Dfd* in *O. sagittarius* and *O. taurus* recapitulated the ventral homeotic transformations that resulted from the double knockdown treatment (Figure 4H, Figure S1D). Collectively, the nature and extent of mouthpart transformations observed here thus largely match those reported previously in *Tribolium* beetles (Smith & Jockusch 2014), validate our RNAi methodology in dung beetles, and indicate a high degree of conservation of *Hox* patterning function in the ventral head during the metamorphic transition in Coleoptera.

Likewise consistent with our initial predictions, two of our three *Hox* gene knockdowns failed to alter *dorsal* head formation. Specifically, *Os-pb* and *Ot-pb* RNAi animals did not exhibit any dorsal head patterning alterations within or outside horn-bearing regions or the corresponding hornless regions in alternate sexes (Figure S2A). Similarly, *Os-Dfd* and *Ot-Dfd RNAi* animals did not exhibit any dorsal head patterning defects (Figure S2B), nor did animals obtained from the double knockdown treatment (Figure S2C).

However, *lab* RNAi did result in an unexpected dorsal head phenotype in both *O. taurus* and *O. sagittarius*. In particular, while *lab* RNAi animals did not reveal any obvious alterations with respect to horn presence, position, size, or shape (Figure 5A-B), *lab* RNAi treatment did result in an unexpected bilateral patterning defect in the posterior-most dorsal head of both species. Specifically, *lab* RNAi resulted in a deformation of the *tempora* (or *temples*; *sensu* Steinman & Zombori 2012, Lawrence & Slipinski 2013), which in wildtype and control-injected individuals represent the typically conspicuously everted dorsal postero-lateral corners of the head located behind the eyes. *lab* RNAi individuals, in contrast, lacked everted tempora in males and females of both species, resulting instead in a conspicuous narrowing of the dorsal posterior-most head (Figure 5C-D).

**Figure 5.**
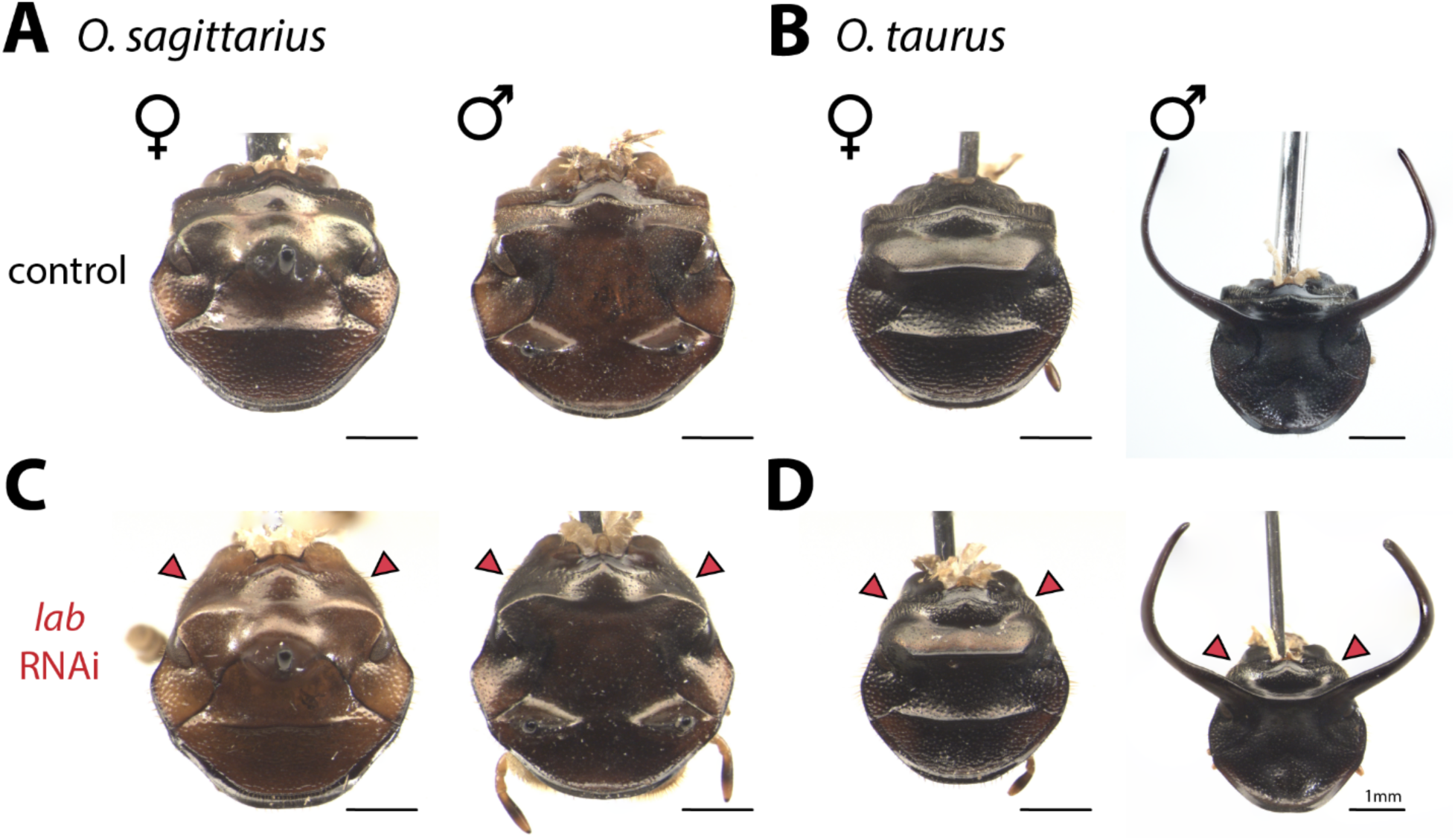
*lab* RNAi results in the loss of the tempora located in the dorsal posterior head. Dissected heads of buffer-injected control beetles illustrating normal dorsal head morphology for *O. sagittarius* (A) and *O. taurus* (B). *Os-lab* RNAi did not affect horn presence, positioning, size, or shape in male or female adults, but did result in the bilateral deformation of the posterolateral tempora (temples) in both sexes (C). *Ot-lab* RNAi similarly did not affect horn presence, positioning, size, or shape in males, but again resulted in bilateral deformation of the temples in both sexes (D).

Lastly, triple knockdown of *lab+pb+Dfd in O. sagittarius* resulted in a combination of dorsal and ventral phenotypes seen in single gene RNAi treatments, with animals exhibiting the bilateral reduction and rounding of temples in addition to mouthpart transformations (Figure S2D). In partial contrast, triple knockdown of *lab+pb+Dfd* in *O. tauru*s only recapitulated the ventral homeotic transformations, and did not result in animals exhibiting rounded corners in the dorsal posterior head (Figure S2E).

### Ablation-based fate mapping confirms correspondence between labial expression in larvae and adult labRNAi phenotypes

To further establish a link between the juvenile expression of *Os-lab* in the presumptive intercalary segment to the loss of adult temples following *lab* RNAi, we ablated head epithelial cells in *O. taurus* and *O. sagittarius* larvae prior to their metamorphic transition (as in Busey *et al*. 2016). We focused on three separate regions of the posterior dorsal head, targeting cells on the right side of the head of each treated larvae and leaving the left side of the head untreated to serve as a negative control (Figure 6A, Supplementary Table 3). Note that for ease of viewing of the temple region, we took advantage of the high nutrition sensitivity of horn development in *O. taurus* to rear smaller-sized males, such that their correspondingly smaller head horns would not obscure viewing of the *tempora* from a dorsal perspective.

**Figure 6.**
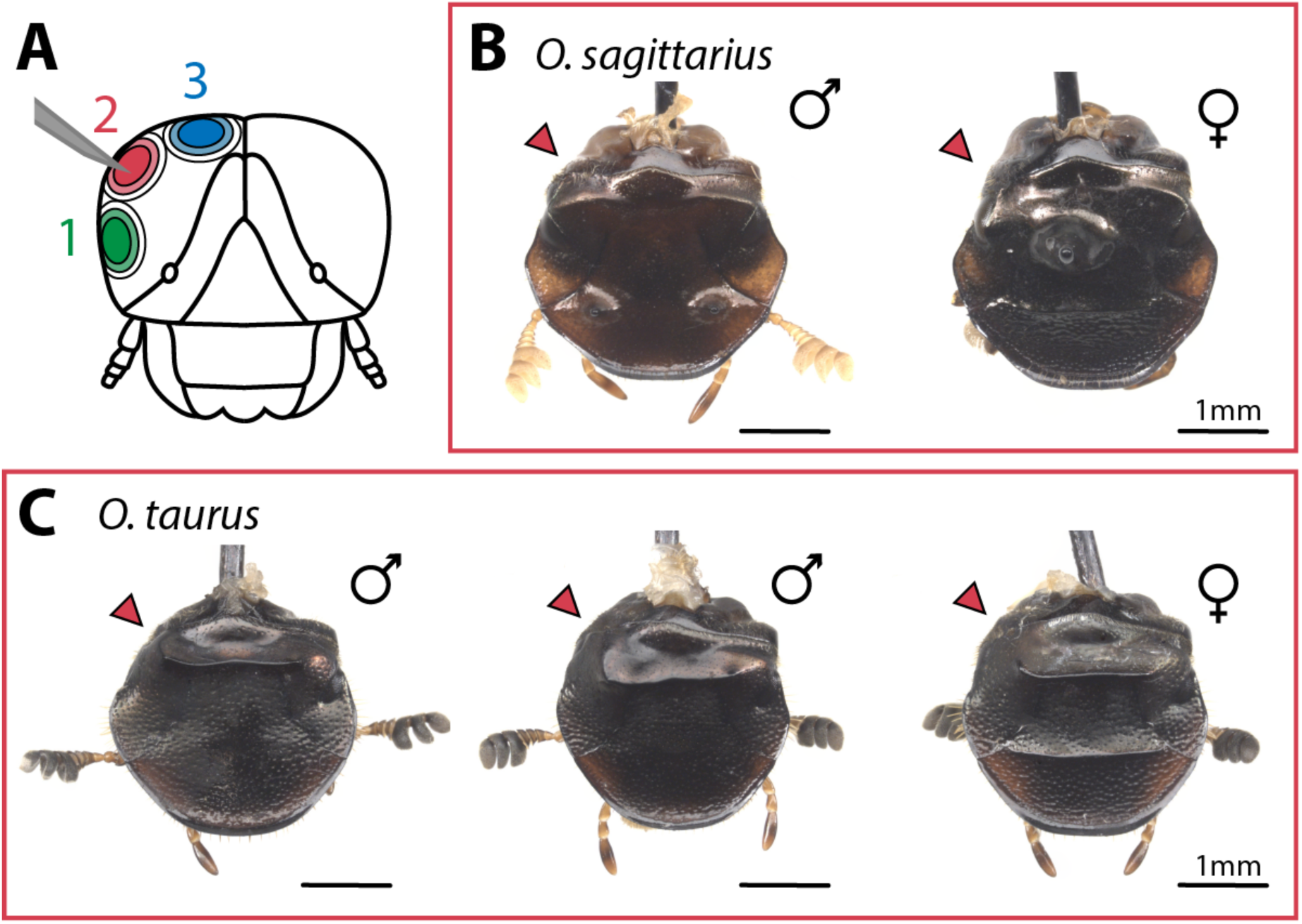
Ablation-based fate mapping of the larval head recapitulates loss of temples seen in *labial* RNAi phenotypes. Diagram of the larval head regions that were unilaterally targeted through ablation-based fate mapping (A). Ablation of region 2 resulted in a flattening of the posterolateral tempus, phenocopying the result of *lab* RNAi in *O. sagittarius* adults (B) and *O. taurus* adults (C; two representative individuals are shown). Phenotypes resulting from ablation of regions 1 and 3 can be found in Supplementary Figure 3.

Ablation targeting region 1, the most lateral region, resulted in unilateral loss of the right compound eye in both species, while targeting region 3, closest to the posterior dorsal midline of the head, resulted in no observable morphological defects (Figure S3). Ablation targeting region 2 resulted in unilateral rounding or flattening of the normally everted right temple of the cuticular vertex of the posterior head in both species, thus phenocopying *lab* RNAi results. Note that ablation of region 2 in *O. taurus* (but not the larger *O. sagittarius*) also consistently eliminated the right adult compound eye. This suggests that in this species the larval intercalary epithelium and the larval ocular primordia are in such close spatial proximity that the relatively coarse-grained ablation approach taken here could not target region 2 in *O. taurus* without also affecting region 3 due to the much smaller size of *O. taurus* larvae.

## Discussion

Motivated by preliminary data documenting unexpected *Hox* gene expression in the dorsal heads of beetle larvae, we investigated the genomic *Hox* content and arrangement in multiple species of horned dung beetles. We then assessed embryonic *Hox* expression patterns and tested for the potential function of anterior *Hox* genes *labial, proboscipedia,* and *Deformed* in establishing the segmental boundaries of the adult beetle head and in patterning their highly diversified, evolutionarily novel cephalic horns. Below we discuss the most important implications of our results for our understanding of the development, evolution, and diversification of the insect head.

### Onthophagus sagittarius labial is translocated within the Hox cluster yet exhibits conserved intercalary segment expression

Our reciprocal BLAST approach followed by secondary validation of cluster contiguity on PacBio sequences indicated that the canonical insect *Hox* cluster is intact with a full complement of genes in two of the three species investigated, *O. taurus* and *D. gazella*. In partial contrast, while a similarly complete complement of Hox genes present on a single scaffold was uncovered in *O. sagittarius*, the cluster arrangement in this species was unexpected, with the majority of the 3’ cluster (from *Os-pb* to *Os-abdB*) translocated ∼8MB past the 5’ end of the cluster nearing the end of the scaffold, with the most 5’ gene *Os-lab* alone maintaining its more central scaffold location (Figure 2). The conditions and mechanisms that may have led to this translocation remain unclear, however, recent comparative genomic analyses of this clade have uncovered a massive degree of transposable element expansion across the *O. sagittarius* genome (Davidson & Moczek 2023), indicating that transposable element insertions and subsequent inversions could be one potential mechanism underlying this cluster rearrangement in this species.

While the *Hox* gene cluster is known for its deeply conserved collinearity, cluster splits, inversions, and gene translocations are not wholly uncommon (Lemons & McGinnis 2006, Mulhair & Holland 2022). For example, a study of seven species within the genus *Drosophila* established that none of the species exhibited the single closely linked complex that is putatively ancestral for the genus and seen in other Diptera, each of them instead possessing one or more major splits (Negre & Ruiz 2007). Specifically, two of the major cluster splits were dated to have occurred between 63 and 43 million years ago, with one more recent split having occurred between 30 and 20 million years ago. Other taxa such as the tardigrades have sustained major *Hox* gene loss, retaining only five of the eight body patterning genes (Smith et al 2016). Our results here contribute to this body of work and document an exceptionally recent cluster translocation, with phylogenetic estimates dating the most recent common ancestor of *O.sagittarius* and *O. taurus* to within the last 5 million years (Davidson & Moczek 2024).

Furthermore, experimental manipulations have shown that presence of enhancer sequences within the endogenous cluster orientation are crucial for proper regulation of Hox gene function (Tschopp & Duboule 2011, Tschopp *et al*. 2011). Experimentally generated inversions and translocations separating normally linked enhancers from their target Hox genes phenocopy deletions of those coding sequences, indicating that evolutionary separation of a Hox gene from its ancestral regulatory landscape may be sufficient to alter its function.

Given that the derived cluster arrangement in *O. sagittarius* specifically affected *labial,* i.e. the *Hox* gene expressed most anteriorly, and that this species possesses a highly derived head horn morphology relative to the ancestral state for this clade (Emlen *et al*. 2005), we next pursued expression and function analyses of the anterior-most Hox genes *labial* and *proboscipedia* to assess the possible functional significance of this derived cluster configuration. Specifically, we sought to assess whether the genomic rearrangement could underlie a derived function in regulating the patterning of the *O. sagittarius* head and/or the cephalic horns contained therein. Using *in situ* hybridization chain reaction, we found *lab* and *pb* expression patterns to be conserved: specifically, we detected *Os-labial* expression during early embryogenesis within the presumptive intercalary segment posterior to the non-segmental ocular region, whereas in later embryos *Os-lab* expression was observed anterior to the mouthparts and restricted to two distinct lateral regions of the head (Figure 3F-H). *Os-proboscipedia* in turn was expressed – again following expectations – in the segments directly posterior to the region marked by *lab* expression, bearing the maxillary and labial mouthparts (Figure 2).

### RNAi targeting anterior Hox genes transforms mouthparts as expected but does not alter head horn formation

Famously, Hox gene manipulations result in homeotic body segment transformations, including the corresponding appendages generated by each segment (Carroll 1995). The original studies in *Drosophila* revealed a functional hierarchy in the patterning mechanism of the Hox genes termed *posterior prevalence.* Functional consequences of this phenomenon include ectopic expression of a posterior Hox gene in an anterior segment overriding normal segment identity, and the attenuation of posterior Hox expression allowing more anterior segment identities to become de-repressed (Lewis 1978, Yao *et al*. 1999). This phenomenon is controlled on a molecular level by the hierarchical dominance of posterior Hox genes over the function of more anterior Hox genes, and is achieved by a diverse set of mechanisms including mutual transcriptional repression between co-expressed Hox genes (Hafen *et al*. 1984, Carroll *et al*. 1986).

Data from embryonic studies in *T. castaneum* established that Hox genes generally repress antennal identity throughout the more posterior gnathos and trunk (Brown *et al*. 2002). More recently, Smith & Jockusch (2014) assessed the roles of *Hox* identity specification during metamorphosis and documented the mouthpart transformations that occur in adult *T. castaneum* after larval *Hox* RNAi. These data led us to predict that *Hox* manipulations knocking down *pb* and *Dfd* in *Onthophagus* beetles would result in a matching set of homeotic transformations of the mouthparts; in contrast, manipulations targeting *lab*, which specifies the appendage-less intercalary segment, have not been documented to cause homeotic phenotypes in adults (Smith & Jockusch 2014).

Our data closely matched these prior results (Figure 4). RNAi targeting *lab* did not generate any mouthpart transformations, as expected, while attenuation of *pb* expression resulted in the transformation of two gnathal appendages, the maxillary and labial palps, into legs. Based on the logic of posterior prevalence, this phenotype is explained by the anterior migration of Scr and Antp expression, two genes which normally regulate appendage identity in the thorax, now specifying thoracic identity in gnathal segments (Abzhanov *et al*. 2001). Attenuation of *Dfd* expression in turn resulted in deformations of the base of the maxilla and mandible, but not in homeotic transformations. This phenotype is likely due to the maintenance of gnathal segment identity by *pb* expression that is retained in the gnathal segments (Brown *et al*. 2002). Double knockdown of *pb* and *Dfd* resulted in the transformation once again of labial palps to legs, but of maxillary palps into antennae; this is likely due in part to anterior migration of *Scr* and *Antp*, along with new posterior expression of antennal specification genes in the anterior head, in line with findings in other insects illustrating that antennal identity is the default state that will be expressed in absence of any *Hox* gene expression (Brown *et al*. 2002). Taken together, our suite of results provides support for a high degree of conservation across Coleoptera for the regulation of gnathal segment patterning by *Hox* genes during metamorphosis.

In addition to assessing these ventral mouthpart transformations, we sought to test whether *Hox* RNAi might also result in morphological effects on the cephalic horns in the dorsal head due to the detectable dorsal expression of these genes during the early stages of metamorphosis (Linz and Moczek 2020) and the previously unknown boundaries of the intercalary segment in the post-metamorphic head. However, larval *Hox* RNAi treatments did not result in any cephalic horn patterning defects, in either the species with canonical or derived *Hox* cluster arrangements. Across multiple distinct horn morphologies (paired lateral posterior horns in male *O. taurus*, a single medial posterior horn in female *O. sagittarius*, and paired anterior horns in male *O. sagittarius)*, *Hox* RNAi treatments did not affect horn presence, positioning, number, size or shape. Instead, these results are consistent with earlier ablation fate mapping findings (Busey *et al*. 2016) that suggested that the diversity of cephalic horns found in *O. taurus* and *O. sagittarius* do in fact derive solely from the non-segmental ocular region of the head rather than any segmentally derived tissues.

### Labial knockdown and ablation-based fate mapping approaches resolve beetle head segment boundaries through metamorphosis

Even though our RNAi treatments targeting *lab* did not affect horn forming regions, it did nevertheless reveal an unexpected morphological patterning deffect in the dorsal head outside horn forming regions. In *Os-lab* and *Ot-lab* RNAi treatments, males and females of both species exhibited a bilateral reduction and rounding of the posterolateral temples, which are positioned behind the compound eyes and just anterior to the head’s articulation with the prothorax (Figure 5). Importantly, these regions qualitatively correspond to the *Os-lab* expression pattern observed in the late embryonic stages (Figure 3), as well as the strong lateral expression detected in transcriptomic studies at the prepupal stage in *O. taurus* (Linz & Moczek 2020). Furthermore, these results are qualitatively congruent with a regional deletion reported in *T. castaneum* larvae following parental RNAi (Posnien & Bucher 2010). However, previous studies in *T. castaneum* failed to find any effect of larval *lab* RNAi on *adult* head structure (Smith & Jockusch 2014). Therefore, to further asses the relationship between juvenile *lab* expression data to the *lab*RNAi phenotypes observed in *Onthophagus* adults, we performed ablation-based fate mapping during larval development. The fate mapping data confirmed that ablation of a targeted region of the posterolateral larval head affects the same posterolateral region of the adult head in ways similar to those induced by larval *lab* RNAi (Figure 6). Taken together, our work has generated data that begin to establish correspondence between embryonic, juvenile, and adult head segments of onthophagine beetles (Figure 7). These findings have a range of possible implications, which we will discuss in turn.

**Figure 7.**
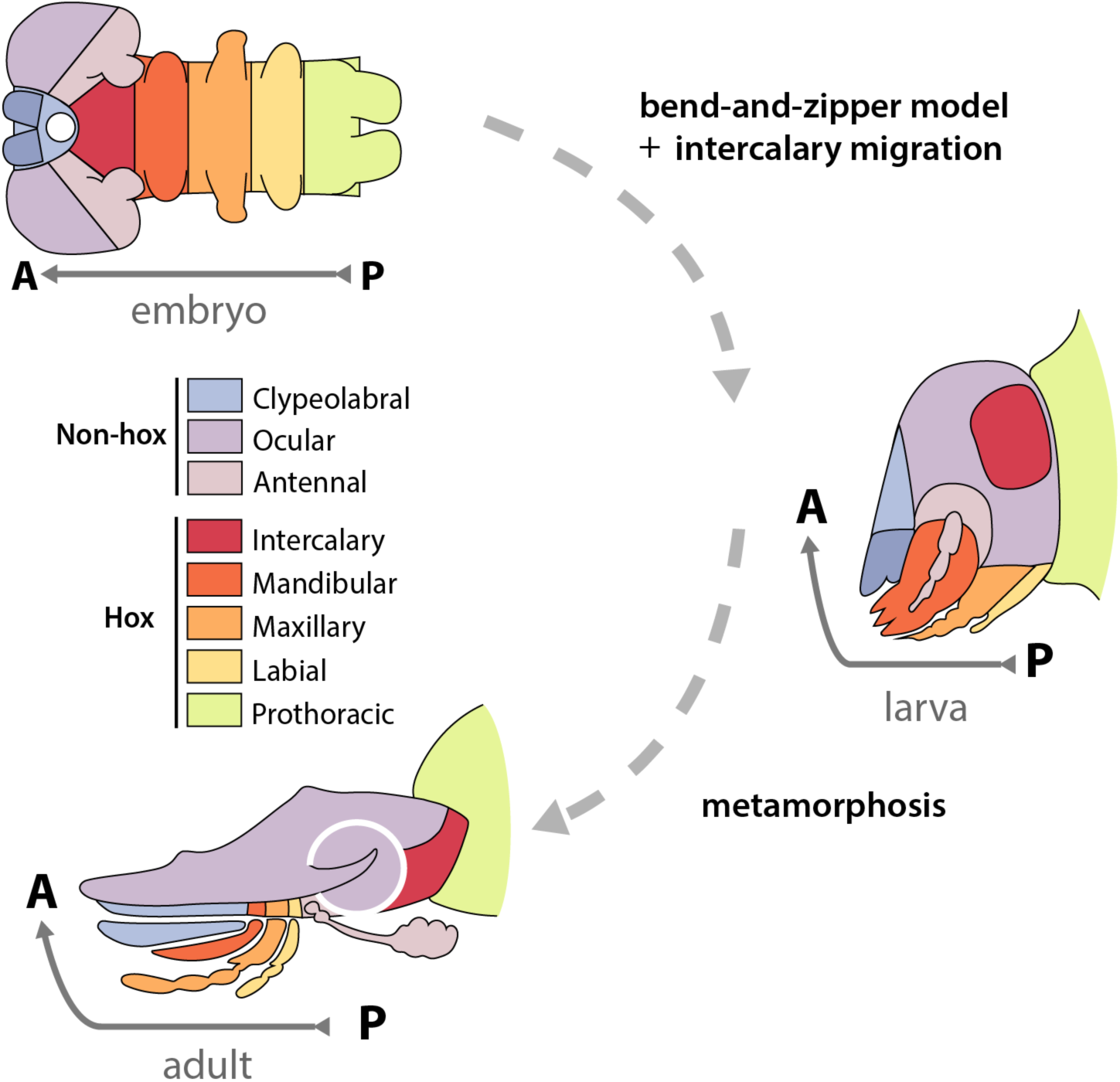
Pregnathal segment boundaries throughout Onthophagine development. *lab* expression data, RNAi phenotypes, and ablation fate mapping results suggest that the posterolateral temples of the vertex of the adult beetle head derive from the intercalary segment.

One possible interpretation of these data is that these phenotypes continue to reflect a reliable fate map for the intercalary segment throughout embryonic, juvenile, and adult stages of *Onthophagus* head morphogenesis. If so, this interpretation would imply that the anterior-dorsal closure motion proposed by the bend-and-zipper model may hold for the majority - but not entirety - of the dorsal *Onthophagus* head. Rather than extending fully across the mediolateral axis covering the intercalary, maxillary, mandibular, and labial segments positioned on the ventral side of the head, as proposed earlier, our findings raise the possibility that the dorsal closure of the ocular region may extend all the way to the prothoracic segment only in the medial head. If so, the bilateral adult *Onthophagus* genae may derive instead from the intercalary segment that has separated and migrated bilaterally to form the posterolateral head, as such matching data reported in larval *Tribolium* by Posnien and Bucher (2010). While this interpretation would seem at odds with a lack of phenotype reported in adult *Tribolium* following larval *lab*RNAi, this discrepancy could be accounted for by the fact that the morphology of the adult *Tribolium* gena is more flattened, simplified by comparison, and does not generate any protrusions. Thus any effect of *lab*RNAi may have been too subtle to detect, as neighboring head epithelial cells may have been able to proliferate and compensate for the deletion of the intercalary segment, producing a morphology roughly equivalent to that of wildtype in the process.

A similar scenario that we cannot reject based on current data is that the intercalary *Onthophagus* segment bifurcates and migrates bilaterally within the head, yet like in adult *Tribolium* does *not* contribute to the externally visible genae. The flattening of the temples seen after *lab*RNAi could then be explained by the deletion of internal tissue that no longer scaffolds everted temples. This interpretation would again account for the lack of corresponding RNAi phenotypes found in adult *Tribolium*.

Alternatively, we cannot fully exclude the hypothesis that evolutionary changes in the timing and/or spatial distribution of *lab* expression and function among beetle families may account for differential presence and absence of adult labRNAi phenotypes. For example, in *Tribolium* larvae the segmental contributions to the head include a lateral contribution of the intercalary segment to the genae. During metamorphosis the ocular and/or antennal regions appear to be replacing this contribution in *Tribolium,* but may not do so in *Onthophagus,* thereby explaining the absence and presence of adult *lab*RNAi phenotypes, respectively.

While the above scenario posits an evolutionary change in labial function across developmental *time*, changes to labial’s *spatial* expression may similarly acount for the results presented here. For instance, the domain of *lab* expression may have expanded in *Onthophagus* relative to *Tribolium*, yielding a larger internal and/or external head contribution in the process. Given that the molecular mechanism of mutual transcriptional repression between Hox genes does not limit expansion of labial into the ocular region, this scenario cannot be ruled out.

While data at present do not allow us to distinguish between these alternative scenarios described above, collectively, our results document that *labial* functions in patterning the dorsal head in beetles, a region previously thought to be derived from the non-segmental *Hox-free* pre-ocular region of the embryo. These data allow for reconciliation of the unexpected *lab* expression data detected in previous work (Linz & Moczek 2020), by shedding light on more complex processes of adult head morphogenesis in the derived, flattened Onthophagine head morphology. However, this work also established a lack of functional consequences for *pb* RNAi in the dorsal head, indicating that this expression data detected in the dorsal head must be latent, or was derived from more internal tissues not reflective of the segmental origin of the local epithelia. Furthermore, these results establish a conservation of general *Hox* patterning function despite a completely unexpected genomic rearrangement of the cluster in one of three onthophagine beetles studied and contribute to an understanding of the morphogenetic processes underlying development of the beetle head throughout the life cycle.

In studies on the origin of *prothoracic* horns in beetles, investigation of the Hox gene *Scr* was instrumental in uncovering the serial homology of horns on the prothorax and wings on the meso- and metathorax, among other paired bilateral structures on abdominal segments (Hu *et al*. 2019, Hu & Moczek 2021). Preliminary RNAseq work and discovery of an unexpected rearrangement of the Hox cluster motivated the consideration of anterior Hox genes as potentially relevant in the origin and diversification of cephalic horns. However, the work here documents that Hox regulation has likely not played a role in origin or diversification of cephalic horns, making clear that despite the elegant ecological and behavioral integration of cephalic and prothoracic horns, these two horn types arose through intriguingly distinct evolutionary and developmental mechanisms. Thus, future work must uncover the upstream regulators that enabled the developmental genetic origins of this enigmatic evolutionary novelty.

## Acknowledgements

The authors would like to thank Dr. David Linz for inspiration and assistance with generating figure components, Dr. Eduardo Zattara for assistance with generating the bend-and-zipper model figure, Dr. Phil Davidson for assistance compiling genomic analysis programs, Dr. Frank-Thorsten Krell for assistance in parsing the literature on scarab morphological terminology, and Dr. Andras Kun and the Indiana University Light Microscopy and Imaging Center for providing microscopy expertise and equipment. Comments by Dr. Phil Davidson, Dr. Rebecca Westwick, Joshua Jones, and Kenzie Givens greatly improved earlier drafts. This work was supported in part through generous funding from the National Science Foundation [Grant no. 2243725 and 1901680 to APM] and was performed while EMN was funded by the National Institutes of Health [T32-HD049336]. Additional support was provided by the IU Groups Scholars Summer Research Experience to SG.

## Supplementary Material

**Figure S1.**
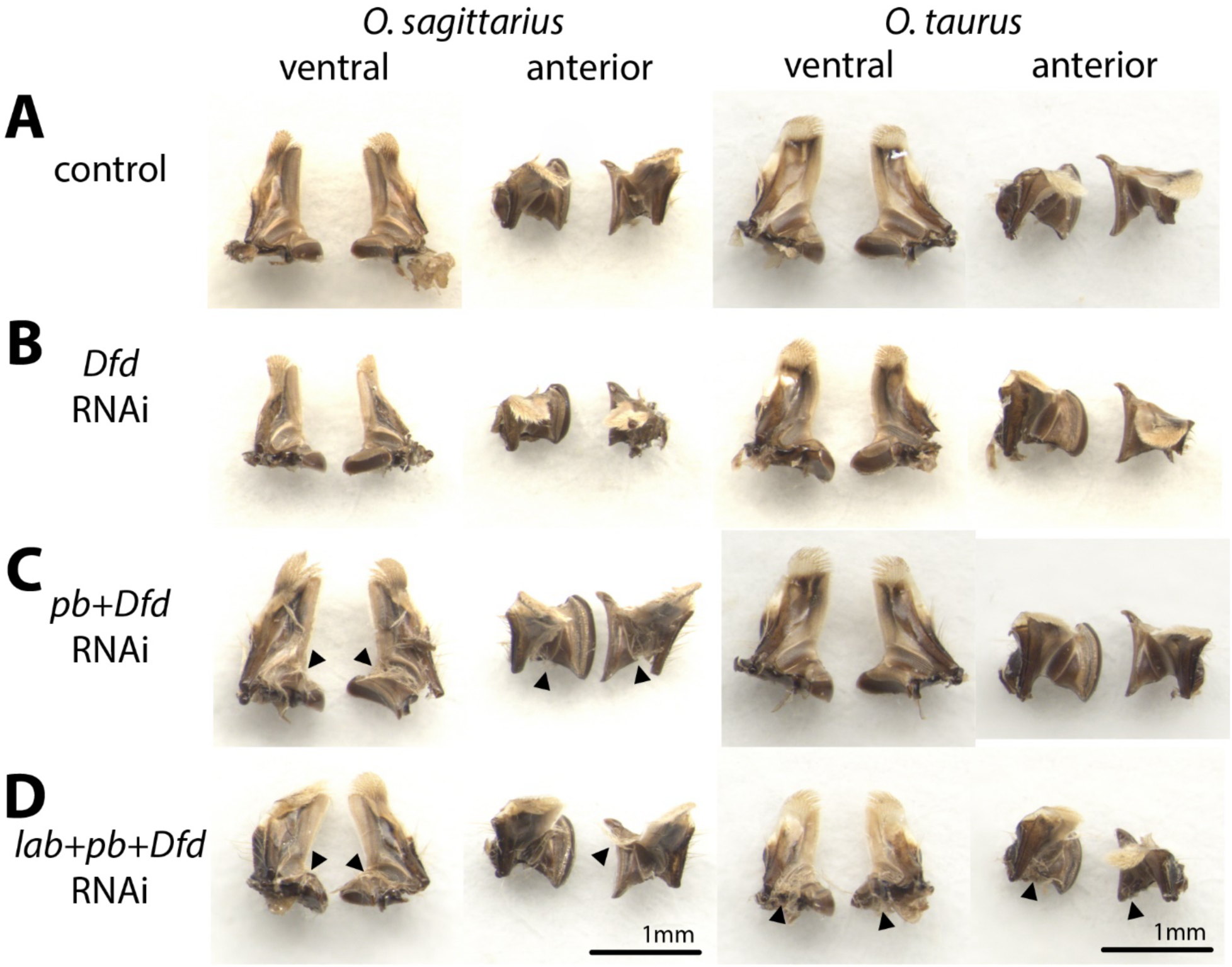
Mandible morphology following RNAi targeting *Dfd,* alone or in combination with *lab* and *pb*. (A) Control-injects *O. sagittarius* and *O. taurus* beetles display wild-type mandible morphology which in onthophagine beetles is specialized for filter feeding on liquids; the proximal end, termed the molar, is smooth and wide, while the distal end, termed the incisor, tapers into a fringed medial side and tip of distal fringe (see ventral view) The proximal mandibles form an asymmetric articulation between left and right sides (see anterior view) (B) *Os-Dfd* and *Ot-Dfd* RNAi appear to have no effect on mandible morphology. (C) RNAi targeting *pb* and *Dfd* did not affect *O. taurus*, but generated ectopic bristles on the molar regions in *O. sagittarius*. (D) RNAi targeting *lab, pb*, and *Dfd* resulted in the appearance of ectopic bristles in both species (arrowheads). Note that the individuals in the left column of B and right column of D were of an overall smaller body size; changes in the shape of the articulation of left and right molar regions is associated with this body size difference.

**Figure S2.**
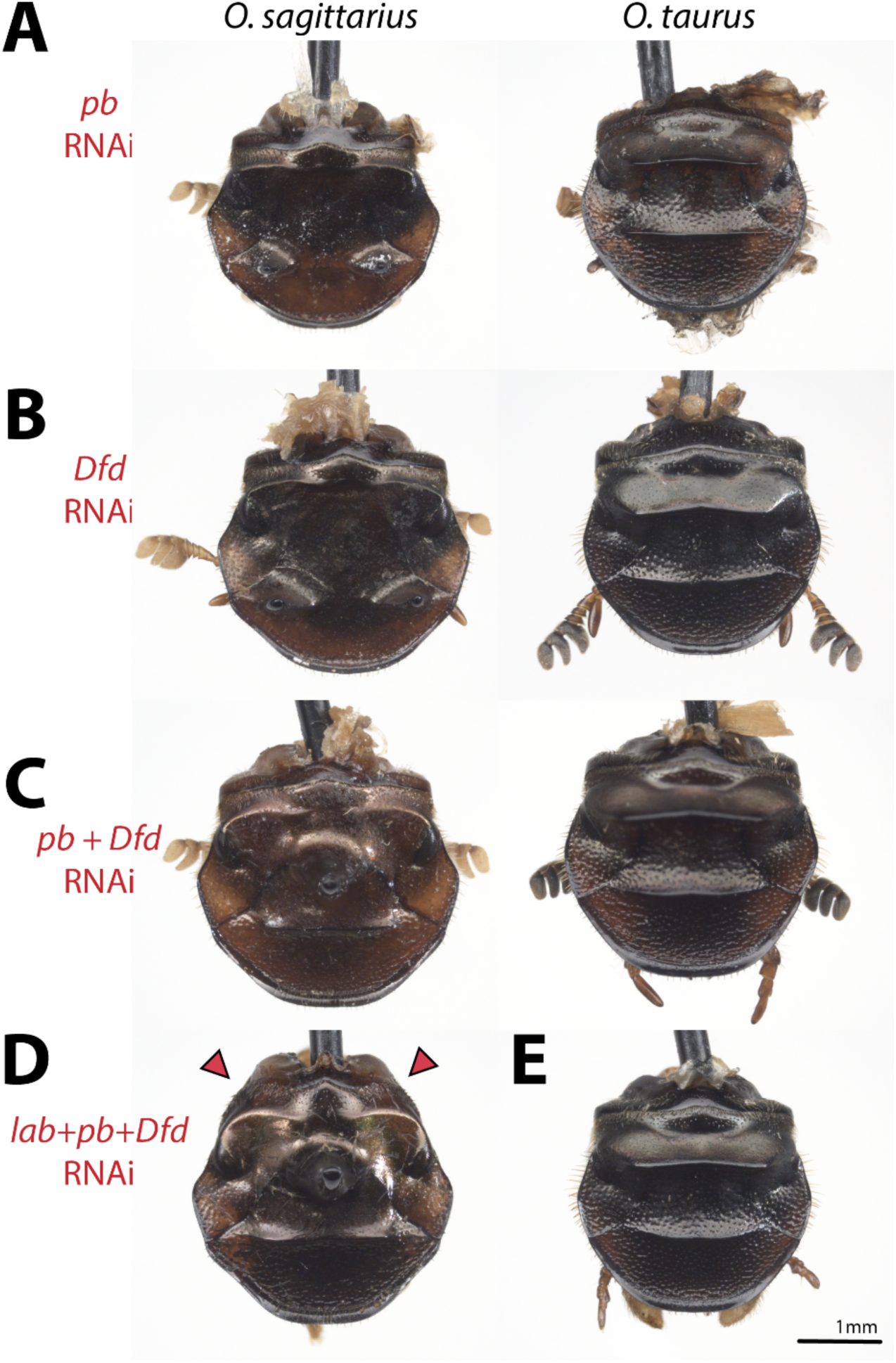
Effect of RNAi targeting *pb, Dfd,* and *lab* alone or in combination on dorsal head formation. Neither (A) *Os-pb* and *Ot-pb* RNAi, (B) *Os-Dfd* and *Ot-Dfd* RNAi, nor (C) RNAi targeting *pb* and *Dfd* simultaneously affected dorsal head formation, including the development of cephalic horns. (D) Triple knockdown of *lab+pb+Dfd in O. sagittarius* recapitulated the flattening of the posterolateral temples of the vertex (D). In contrast, triple knockdown of *lab+pb+Dfd* in *O. tauru*s did not result in any dorsal posterior head defects in the tempus regions or otherwise (E).

**Figure S3.**
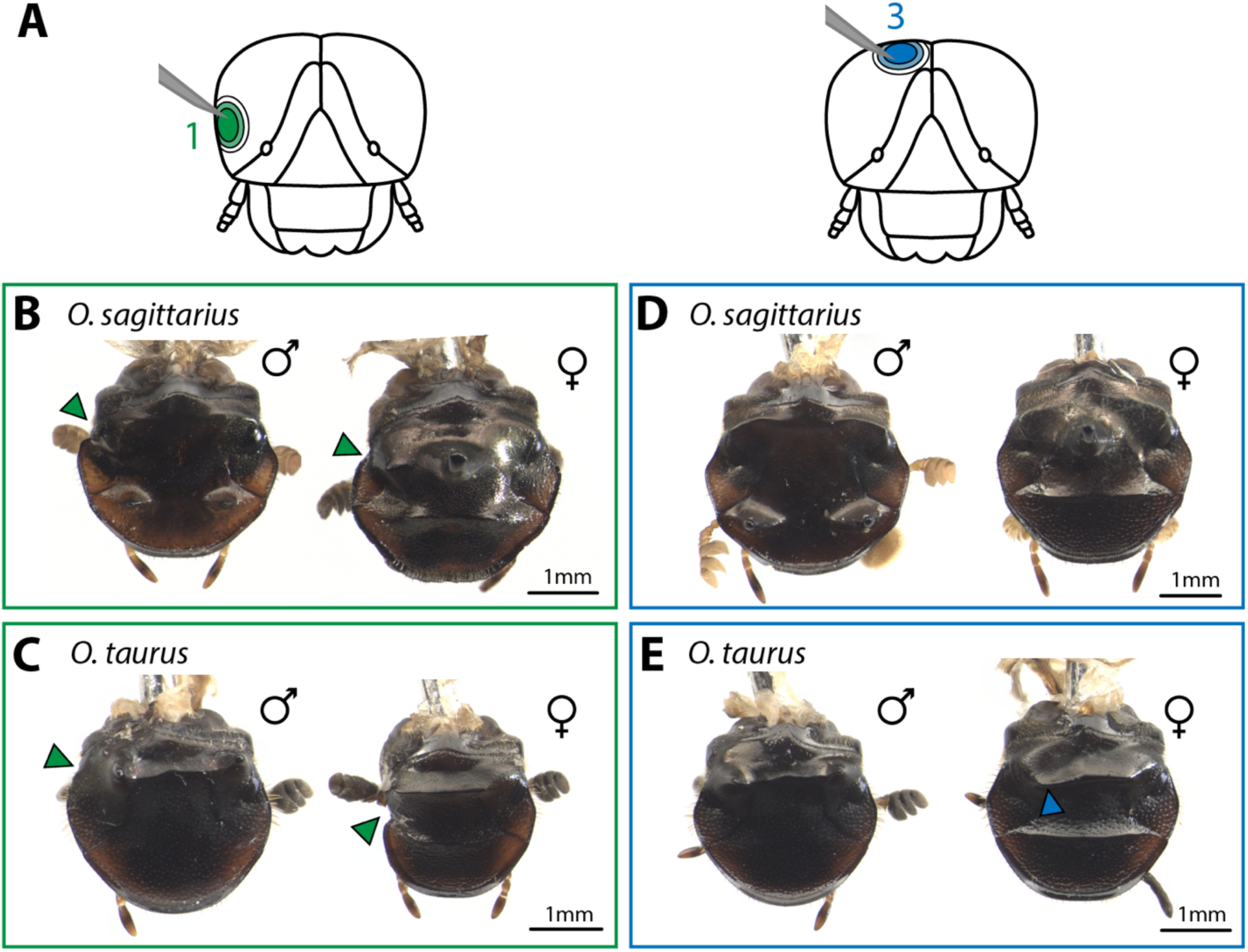
Additional ablation-based fate mapping results. Ablation (A) of head region 1 led to unilateral loss of the right compound eye in (B) *O. sagittarius* and in (C) *O. taurus* (green arrowheads). (D) Ablation of head region 3 yielded no observable morphological defects to the head in most cases (D) with the exception of (E) one *O. taurus* female in which ablation of region 3 disrupted the posterior ridge (blue arrowhead).

**Supplementary Table 1.**
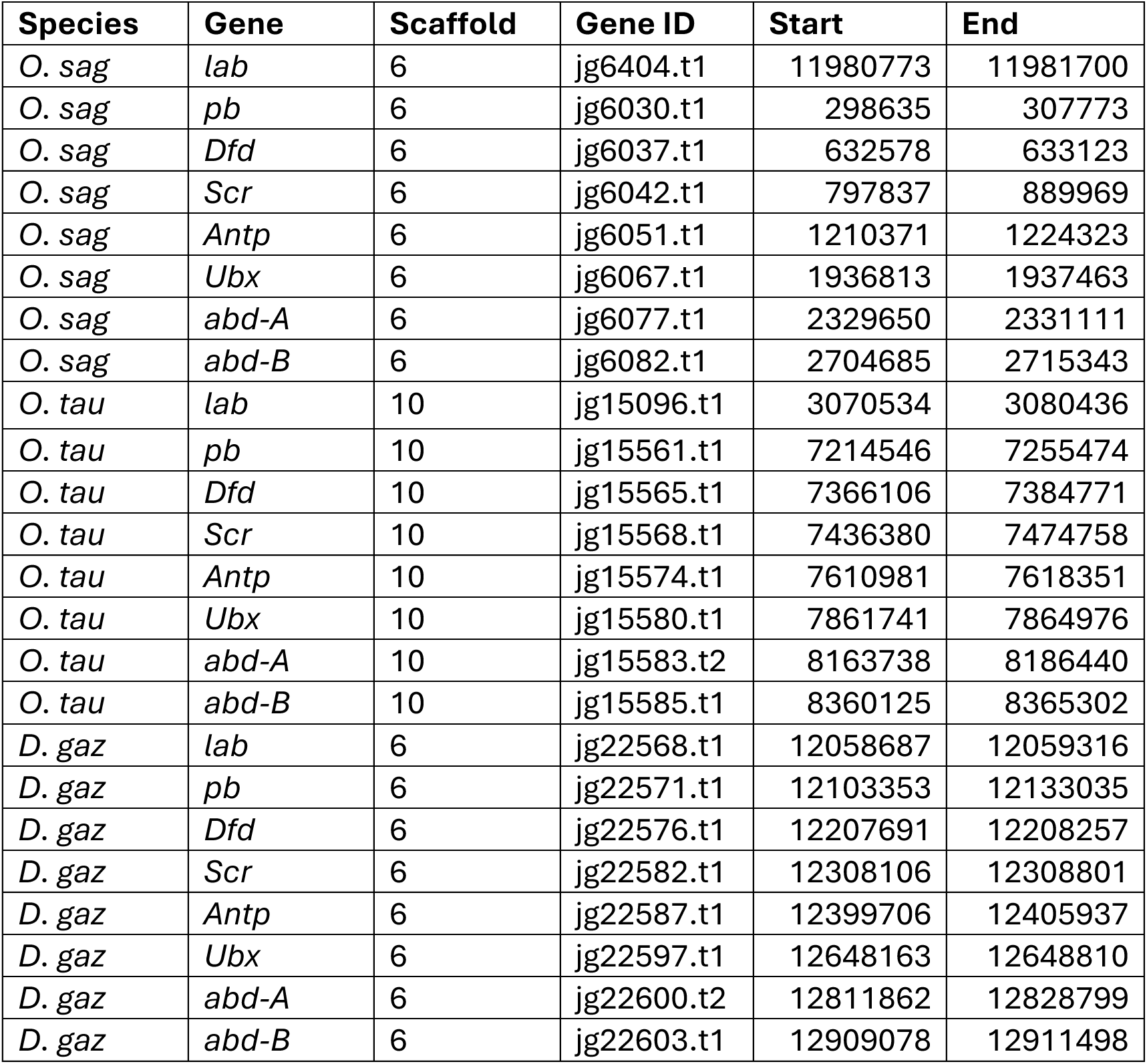
Locations of the *Hox* cluster in Onthophagine genomes.

**Supplementary Table 3.** RNAi and ablation sample sizes and penetrance.

| Treatment | Target | # Treated Larvae | # Dead | # Surviving Adults | # Adult Males | # Adult Females | Penetrance of Phenotype |
| --- | --- | --- | --- | --- | --- | --- | --- |
| RNAi | Os buffer injected | 48 | 7 | 41 | 19 | 22 | - |
| RNAi | *Os-lab* | 72 | 11 | 61 | 28 | 33 | 0.741 |
| RNAi | *Os-pb* | 30 | 14 | 16 | 7 | 9 | 0.625 |
| RNAi | *Os-Dfd* | 24 | 12 | 16 | 11 | 5 | 0.75 |
| RNAi | *Os-pb+Dfd* | 24 | 18 | 6 | 3 | 3 | 1.0 |
| RNAi | *Os-lab+pb+Dfd* | 24 | 19 | 5 | 3 | 2 | 0.8 |
| RNAi | Ot buffer injected | 14 | 3 | 11 | 6 | 5 | - |
| RNAi | *Ot-lab* | 40 | 16 | 24 | 15 | 9 | 0.75 |
| RNAi | *Ot-pb* | 5 | 3 | 2 | 1 | 1 | 1.0 |
| RNAi | *Ot-Dfd* | 24 | 13 | 11 | 7 | 4 | 1.0 |
| RNAi | *Ot-pb+Dfd* | 13 | 8 | 5 | 3 | 2 | 1.0 |
| RNAi | *Ot-lab+pb+Dfd* | 37 | 24 | 13 | 7 | 6 | 0.846 |
| Ablation | Os Region 1 | 25 | 10 | 15 | 7 | 9 | - |
| Ablation | Os Region 2 | 24 | 14 | 10 | 4 | 6 | 0.7 |
| Ablation | Os Region 3 | 19 | 12 | 7 | 5 | 2 | - |
| Ablation | Ot Region 1 | 13 | 11 | 2 | 1 | 1 | - |
| Ablation | Ot Region 2 | 16 | 1 | 15 | 8 | 7 | 0.8125 |
| Ablation | Ot Region 3 | 20 | 3 | 27 | 9 | 8 | - |

